# flowVI: Flow Cytometry Variational Inference

**DOI:** 10.1101/2023.11.10.566661

**Authors:** Kemal Inecik, Adil Meric, Lars König, Fabian J. Theis

**Affiliations:** School of Life Sciences, Technical University of Munich, Freising, Germany 85354; Department of Computer Science, Technical University of Munich, Garching, Germany 85748; Division of Clinical Pharmacology, Ludwig Maximilian University of Munich, Munich, Germany 80337; Department of Mathematics, Technical University of Munich, Garching, Germany 85748

## Abstract

Single-cell flow cytometry stands as a pivotal instrument in both biomedical research and clinical practice, not only offering invaluable insights into cellular phenotypes and functions but also significantly advancing our understanding of various patient states. However, its potential is often constrained by factors such as technical limitations, noise interference, and batch effects, which complicate comparison between flow cytometry experiments and compromise its overall impact. Recent advances in deep representation learning have demonstrated promise in overcoming similar challenges in related fields, particularly in the context of single-cell transcriptomic sequencing data analysis. Here, we propose *flowVI*, a multimodal deep generative model, tailored for integrative analysis of multiple massively parallel cytometry datasets from diverse sources. By effectively modeling noise variances, technical biases, and batch-specific heterogeneity using probabilistic data representation, we demonstrate that flowVI not only excels in the imputation of missing protein markers but also seamlessly integrates data from distinct cytometry panels. FlowVI thus emerges as a potent tool for constructing comprehensive flow cytometry atlases and enhancing the precision of flow cytometry data analyses. The source code for replicating these findings is hosted on *GitHub, ‘theislab/flowVI’*

## 1 Introduction

Modern investigations in immunology demand sophisticated multi-parametric analyses to delineate the elaborate consortia of cell types and perform precise phenotypic categorizations of subcellular states within tissue microenvironments [11, 15]. However, the expansive potential of cellular classifications is still impeded by the stringent technical specifications of flow cytometry method, that allow the investigation of only a limited array of markers at a time, and is heavily dependent on calibration, exhibiting strong batch effects across samples and labs. [3, 5]. While single-cell CITE-seq offers a solution to exceed these constraints, its higher costs, intricate sequencing requirements, extensive antibody panels, and complex experimental protocols deter widespread adoption in the immunological research community [16]. Consequently, there is a pressing need for novel techniques that enable integrated analysis of a multitude of protein expression profiles at the single-cell level, while optimizing accessibility, cost-effectiveness, and high cellular throughput.

Amongst others, this was recently addressed by *Infinity Flow*, IF, a method developed to integrate massively parallel cytometry (MPC) panels (see Figure 1), multiple and partially overlapping datasets to construct a singular, comprehensive data matrix [3]. However, IF is incapable of integrating batches or combining datasets generated from different sources, and necessitates auxiliary methodologies for dimension reduction to investigate cellular heterogeneity. Importantly, the approach also does not decouple the true cytometric signals from background interference, both from non-specific antibody fluorescence and instrument noise, potentially compromising data precision and veracity.

**Figure 1:**
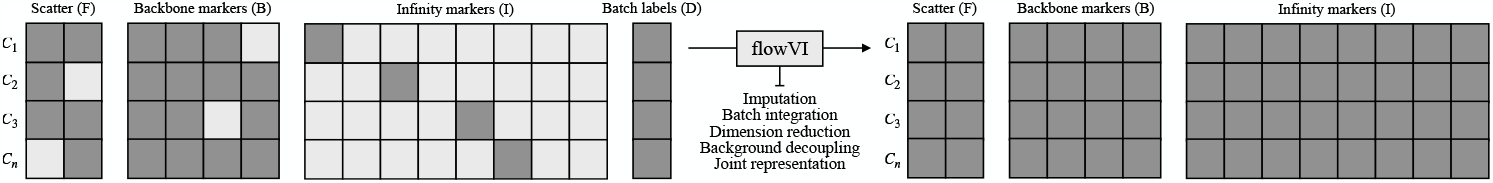
Overview of massively parallel flow cytometry (MPC) analysis with flowVI model. In a typical MPC experiment, as introduced alongside the IF method in Becht et al. [3], each cell is measured for a subset of markers (B) and scatter features (F), and one marker from another subset of markers (I). Unlike IF, where scatter and backbone marker features must be complete, flowVI accommodates missing values and also incorporates batch labels (D) for batch integration tasks.

To address these challenges, we present *flowVI, Flow* Cytometry *V*ariational *I*nference, an end-to-end multimodal deep generative model, designed for the comprehensive analysis of multiple MPC panels from various origins. FlowVI learns a joint probabilistic representation of the multimodal cytometric measurements, marker intensity and light scatter, that effectively captures and adjusts for individual noise variances, technical biases inherent to each modality, and potential batch effects. We show that flowVI’s performance in imputing missing protein markers significantly outperforms the current state of art, the IF method. In contrast to IF, flowVI not only excels in diverse analytical tasks, such as dimensionality reduction for cell-type clustering and background normalization of input features, but also pioneers the integration of multiple MPC datasets, even with limited overlapping panels. This positions flowVI as a promising method for constructing expansive single-cell flow cytometry atlases, while also supporting and standardizing flow cytometry data analysis.

## 2 Methods

Flow cytometry data has a specific structure characterized by disparate distributions observed for scatter features and marker features, each offering unique insights into cellular attributes [1]. This multifaceted measurement necessitates a multimodal analytical framework to comprehensively encapsulate the holistic cellular information. Additionally, the necessity to address the intricacies of flow cytometry data, especially its inherent sparsity, noise, and variability led us to the formulation of flowVI, conceptualized specifically as a multimodal conditional variational autoencoder (cVAE). VAEs are widely acknowledged for their adeptness at modeling complex data distributions and their capacity to encode such data into a meaningful latent space [2, 4, 9]. This capability ensures the preservation of vital information and streamlines various downstream analyses, including clustering, data imputation, trajectory inference, label transfer, and batch integration [8, 10, 13].

To address these intricacies, it is crucial to understand the distinct structures of the input data, namely marker and scatter features. Marker features are continuous and have positive support, reflecting the nature of fluorescence intensity of an antibody-based assay. This data is conceptualized as comprising two components: the foreground signal, representing the abundance of a protein of interest, and the background noise, arising from nonspecific antibody bindings and instrumental errors. Similarly, scatter features, which are also continuous with real support, comprise two factors: the foreground signal, indicative of cellular properties like size and granularity, and background noise, primarily attributed to instrumental errors. By adopting a dual-component modeling for each modality, flowVI aims to robustly discern and delineate the essential characteristics of flow cytometry data, thereby enhancing the accuracy and reliability of subsequent analytical outcomes.

Given the observed data sparsity, particularly within the infinity markers panel as shown in Figure 1, both the inference and generative models are provided with one-hot encoded vectors, *c*_1_, that encapsulate the input data’s sparsity characteristics. Concurrently, both models are informed about batch labels as one-hot encoded vectors, (*c*_2_), during dataset integration tasks. Batch labels are vital for differentiating global founder effects from real biological variations in cross-comparing multiple MPC experiments, like contrasting diseased and healthy samples, thereby enhancing the robustness and relevance of the analysis. Joint input feature vectors, constructed by concatenating observed data from marker (*x*_*m*_) and scatter (*x*_*s*_) modalities, are fed into a shared inference module. The joint posterior is then sampled via the reparameterization trick [12], to obtain the joint embedding, *z*_joint_.

Joint latent space representations then are channeled into the modality-specific decoders reconstructing the corresponding input data. For the scatter features, three of neural networks modules are employed to reconstruct the observed data under Gaussian mixture loss function, Equation 1 and 3, where the distribution is specified by the mean *μ*_1,2_, standard deviation *σ*_1,2_ and mixing coefficient *π*. Simultaneously, the marker data is decoded through four neural network modules, leveraging a Gamma mixture loss function, Equation 2 and 3, where shape *α*_1,2_, scale *β*_1,2_, and mixing coefficient *π* characterize the distribution. A Kullback-Leibler (KL) divergence term is calculated under the assumption of a Normal distribution in the latent space, Equation 4, where *μ*_joint_ and *σ*_joint_ estimated by the inference model. Additionally, an auxiliary KL divergence term, as described in Equation 5 [7], is implemented to regularize the marker-specific prior for the background mean.

To improve the performance of the generative model in counterfactual imputation, carried out through adapting the input sparsity one-hot vector *c*_1_ during prediction time, a discriminator module is implemented. A cross entropy loss function, Equation 6, on discriminator module predictions enables inference model to minimize the distribution discrepancy between the transformed and target samples. Incorporating the discriminator module aims to create a latent space devoid of well-specific information from the underlying MPC experimental setup, focusing instead on higher-level representations of cellular states regardless of the source well. This strategic implementation ensures that, within the latent space, the mixture of samples from different wells leads to the formation of clusters that are indicative of the cellular state rather than merely reflecting variations in the measured infinity marker for each well. Such an approach is important in mitigating well-effects and promoting a more accurate and generalizable identification of cell states across varying experimental conditions of a given MPC experiment.

Consequently, the total loss function for the proposed model is described in Equation 7, where *w*_*i*_ are hyperparameters that dictate the relative contributions of each loss component. Model hyperparameters, including *w*_*i*_, underwent an extensive optimization process within a broad hyperparameter space, utilizing a large computer cluster and iterative sweeps. This process led to the selection of a specific set of hyperparameters for each dataset, tailored to maximize the accuracy of counterfactual predictions in subsequent analyses. It is also worth mentioning that the development of flowVI was firmly grounded in the *scvi-tools* framework [6], ensuring flowVI’s compatibility and efficiency within the established workflows of single-cell analysis, thereby improving the user experience.

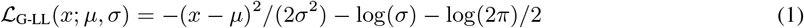

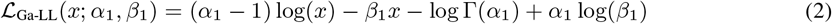

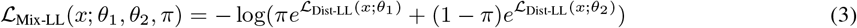

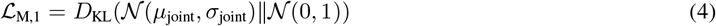

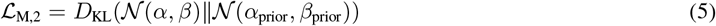

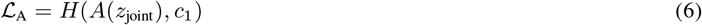

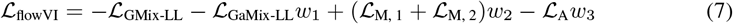

In this study, two large massively parallel cytometry (MPC) datasets were utilized for both model training and subsequent evaluations. The first dataset, *Becht*, sourced from IF method paper [3], comprises 260 infinity markers, 14 Backbone markers, and 6 scatter features. Notably, this dataset doesn’t have specified batch keys for segregating different subsets of data. The second dataset is an in-house unpublished collection, *Konig*, derived from various time points spanning 4 distinct tissues and includes data from 3 individuals. The dataset is characterized by 58 infinity markers, 7 backbone markers, and 6 scatter features, and has 9 batch keys.

## 3 Results

### 3.1 flowVI outperforms Infinity Flow on counterfactual imputations

In assessing the comparative performances of flowVI and IF in imputing missing markers in infinity marker panels, it’s crucial to consider the inherent methodologies in standard flow cytometry analysis.

As discussed by Becht et al. [3] regarding IF performance evaluations, conventional flow cytometry typically employs a progressive gating scheme, discretizing the marker expression levels into distinct bins. Focusing the efforts on the magnitude and frequency of imputed expression, area under the receiver operating characteristic curve (AU-ROC) was thus used as the main metric. In this study, Pearson correlation was also calculated to evaluate the congruence between the predicted and actual values, along with Wasserstein distance to gain insights into the model’s fidelity in mirroring the intrinsic generative distribution.

Our observations highlighted IF’s proficiency in marker imputation, resonating with the insights of Becht et al. [3] for *Becht* dataset. However, flowVI’s imputations displayed superior performance, with an impressive 81% and 97% of the infinity markers better predicted by flowVI for respectively *Konig* and *Becht* (Figure 2a). We observed markers with high AU-ROC values were generally multimodal and frequently expressed, while markers specific to rare cell populations had lower AU-ROCs. Furthermore, in the context of the Pearson correlation of imputed values, flowVI’s prowess is evident, outshining 90% and 97% of the IF predictions respectively. The Wasserstein distance between the two methodologies was found to be statistically indistinguishable.

**Figure 2:**
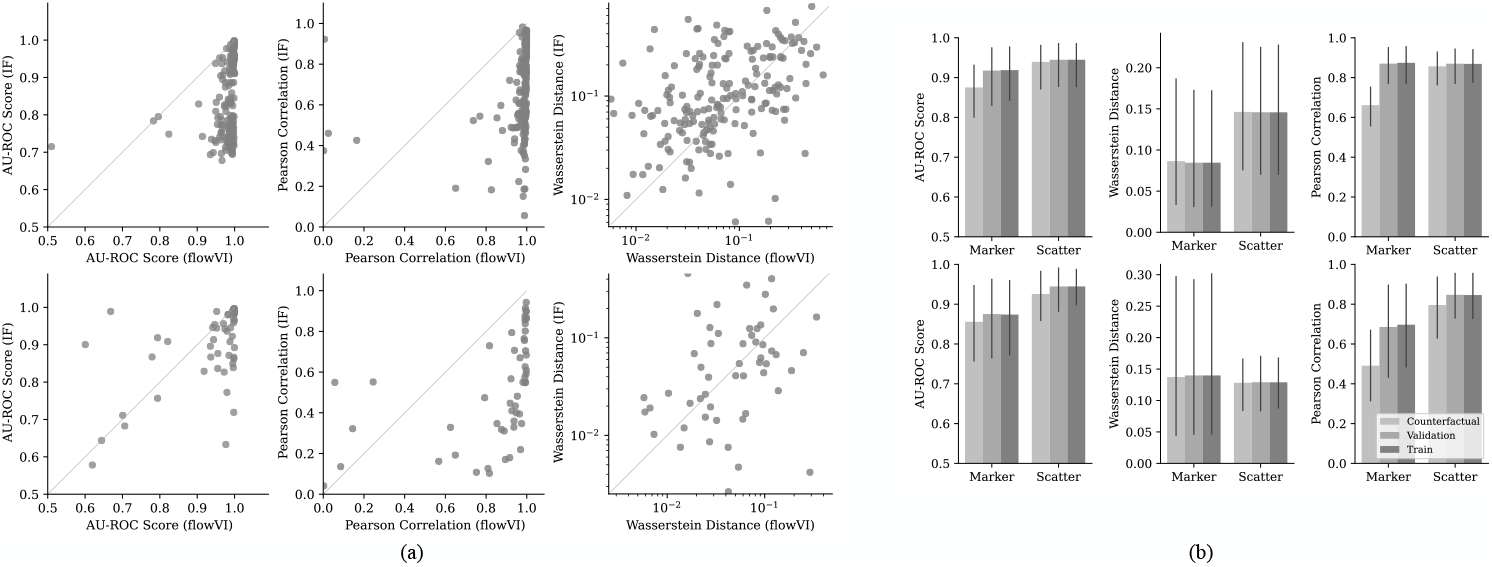
Assessment of model performance and resilience across datasets: *Becht* (upper row) and *Konig* (lower row). (a) flowVI (x-axis) and IF (y-axis) performances on predicting infinity marker expressions on validation sets. (b) Counterfactual imputation through domain adaptation performance of flowVI on noise-added datasets.

While these benchmarks provide invaluable insights, it is imperative to contextualize them within the operational domain of the flowVI model. Notably, flowVI harnesses a domain transfer strategy during its generative phase to ensure robust imputations for unseen infinity marker features, which is carried out through adapting the input sparsity one-hot vector *c*_1_ during prediction time. To evaluate the resilience of the flowVI model under such modified generative conditions, we designed an experiment, where the infinity markers panel was excluded, and each row of the input matrix, comprising both scatter and marker features, had random zero-noise added. Subsequently, we assessed the model’s proficiency in predicting the excluded feature using the retained marker and scatter features. Preliminary observations indicated that while counterfactual imputations marginally underperformed in terms of the main metric, AU-ROC, discrepancies in Pearson scores were more pronounced, suggesting a potential direction for future work to improve the model (Figure 2b).

### 3.2 flowVI effectively integrates multiple massively parallel cytometry (MPC) experiments

The integration of single-cell flow cytometry data, while promising, comes with its own set of complexities and challenges. Integrating single-cell datasets allows for the pooling of data from different experiments, assisting researchers draw conclusions from a broader data spectrum, improving the validity of their findings. It facilitates the comparison of healthy and diseased states on a granular level by consolidating data from different studies, but also strengthens the reproducibility and generalizability of research outcomes. However, it is crucial to approach such integrations with caution, ensuring that the nuances of individual datasets are neither lost nor overshadowed by the collective view.

FlowVI stands out as only proficient method for integrating multiple MPC datasets, even in the presence of limited overlapping panels. Using the limited backbone panel of the *Konig*, a collection 9 different MPC experiments, data points were classified into 5 broad cell-type categories using manual gating strategies. As shown in Figure 3, the flowVI robustly integrates the multiple datasets; mixing different dataset, tissue and samples together, yet astutely clustering identical cell types in the latent space. This indicates that flowVI maintains the integrity of biological information while effectively overcoming the challenges posed by batch differences. Evaluating flowVI’s performance through a battery of metrics for a quantitative understanding, classified into biological conservation and batch integration metrics, flowVI manifested its prowess in seamlessly integrating multiple datasets. In particular, it demonstrated exceptional proficiency in batch correction and attained competitive scores in the preservation of biological nuances (Table 1).

**Table 1:** Performance benchmarks for data integration, obtained by *scib* [14]. Biological conservation metrics assess the preservation of inherent biological signals and essential features such as cell types and states in integrated datasets. Meanwhile, batch correction metrics are designed to benchmark the effectiveness in eliminating batch effects and aligning cells across different datasets, thus ensuring reduced technical variation and enhanced comparability of single-cell data. Here, metrics for the ‘Unintegrated’ are calculated using the combined backbone and scatter panels, while those for flowVI integration are based on latent space embeddings.

| Metric | Bio conservation |  |  |  |  | Batch correction |  |  |  |  |
| --- | --- | --- | --- | --- | --- | --- | --- | --- | --- | --- |
|  | Isolated labels | Leiden NMI | Leiden ARI | Silhouette label | cLISI | Silhouette batch | iLISI | KBET | Graph connectivity | PCR comparison |
| Unintegrated | **0.575** | 0.702 | -11.735 | **0.582** | **1.000** | 0.863 | 0.104 | 0.031 | 0.863 | 0.001 |
| Integrated (flowVI) | 0.559 | **0.752** | **0.069** | 0.565 | 0.998 | **0.901** | **0.228** | **0.114** | **0.902** | **0.803** |

**Figure 3:**
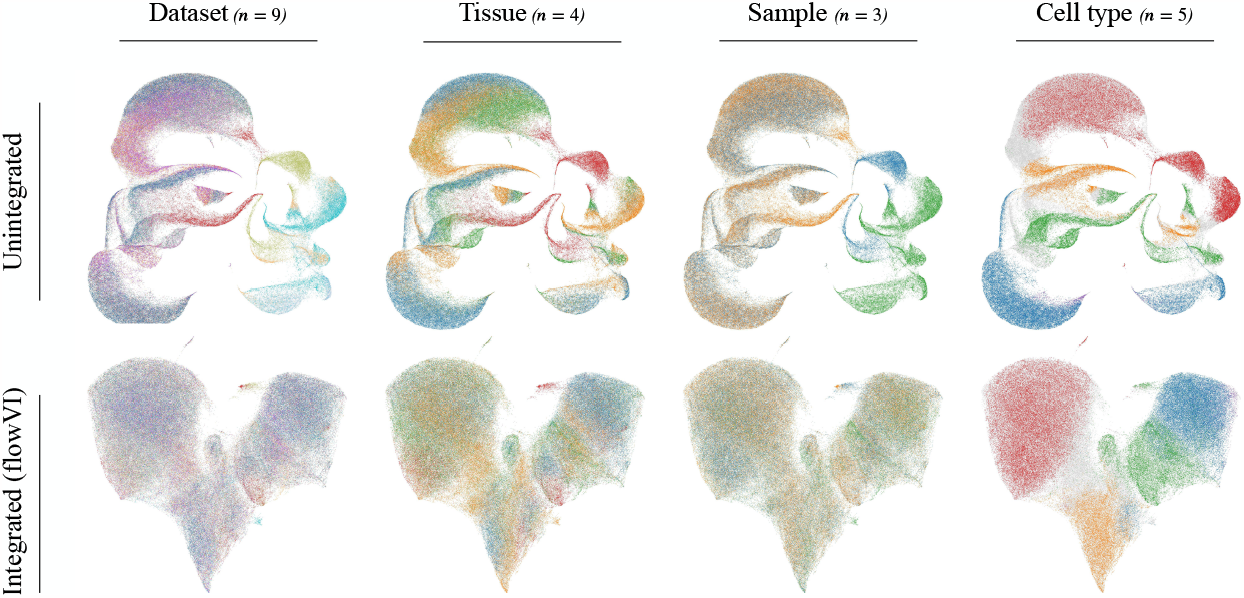
UMAP visualization of multiple MPC experiments with and without flowVI integration for *Konig* dataset. The projection for the unintegrated data utilizes the backbone and scatter panels. In contrast, the flowVI integrated visualization leverages the latent space embeddings.

## 4 Discussion

In this study, we have introduced flowVI as a novel end-to-end deep generative model that excels in the representation learning of multiple flow cytometry datasets. FlowVI accounts for inherent noise and background interference within each modality, leading to an enhanced ability to capture nonlinear relationships between known markers. This enables flowVI to outperform the existing method, IF, for imputing missing markers in massively parallel flow cytometry (MPC) experiments. However, there remains potential for optimizing counterfactual imputations, either through architectural adjustments or by implementing alternative training strategies, in particular by tailoring learning strategies towards domain adaptation for out-of-distribution imputation. Additionally, flowVI effectively integrates multiple MPC datasets from disparate origins, pioneering in this field, thereby promoting comprehensive cross-dataset analyses and advancing our comprehension of pathological conditions. Moreover, in contrast to IF, flowVI generates lower-dimensional latent embeddings suitable for a diverse range of tasks. We envision that flowVI could support the construction of large-scale flow cytometry atlases. This would enable users to engage in cell type identification, including the exploration of rare cell types, within flow cytometry data after mapping to the atlas. In summary, this approach could help move the field beyond the prevalent practice of manual gating for cell type identification and support more standardized cell type harmonization across laboratories.

## Notes

https://github.com/theislab/flowVI

### Competing Interest Statement

F.J.T. consults for Immunai Inc., Singularity Bio B.V., CytoReason Ltd, Cellarity, and Omniscope Ltd, and has ownership interest in Dermagnostix GmbH and Cellarity. The remaining authors declare no competing interests.

